# Crystallographic and kinetic analyses of human IPMK reveal disordered domains modulate ATP binding and kinase activity

**DOI:** 10.1101/434308

**Authors:** Corey D. Seacrist, Raymond D. Blind

**Affiliations:** From the Departments of Pharmacology Division of Diabetes, Endocrinology and Metabolism, Vanderbilt Diabetes Research and Training Center, Vanderbilt University School of Medicine, Nashville, TN 37232; Biochemistry and Medicine, Division of Diabetes, Endocrinology and Metabolism, Vanderbilt Diabetes Research and Training Center, Vanderbilt University School of Medicine, Nashville, TN 37232

**Keywords:** nuclear kinase, kinetics, signaling, inositol phosphate kinase, IPMK, ipk2, PI3-kinase, PIP_2_

## Abstract

Inositol polyphosphate multikinase (IPMK) is a member of the IPK-superfamily of kinases, catalyzing phosphorylation of several soluble inositols and the signaling phospholipid PI(4,5)P_2_ (PIP_2_). IPMK also has critical non-catalytic roles in p53, mTOR/Raptor, TRAF6 and AMPK signaling mediated partly by two disordered domains. Although IPMK non-catalytic functions are well established, it is less clear if the disordered domains are important for IPMK kinase activity or ATP binding. Here, kinetic and structural analyses of an engineered human IPMK lacking all disordered domains (ΔIPMK) are presented. Although the *K*_M_ for PIP_2_ is identical between ΔIPMK and wild type, ΔIPMK has a 1.8-fold increase in *k*_cat_ for PIP_2_, indicating the native IPMK disordered domains decrease IPMK activity *in vitro*. The 2.5 Å crystal structure of ΔIPMK is reported, confirming the conserved ATP-grasp fold. A comparison with other IPK-superfamily structures revealed a putative “ATP-clamp” in the disordered N-terminus, we predicted would stabilize ATP binding. Consistent with this observation, removal of the ATP clamp sequence increases the *K*_M_ for ATP 4.9-fold, indicating the N-terminus enhances ATP binding to IPMK. Together, these structural and kinetic studies suggest in addition to mediating protein-protein interactions, the disordered domains of IPMK impart modulatory capacity to IPMK kinase activity through multiple kinetic mechanisms.

---

As an essential gene in mammalian development with ubiquitous expression in all eukaryotes^1^, IPMK has important functions at the nexus of many important signaling, metabolic and regulatory pathways, exerting influence through both kinase-dependent and independent roles^2^. The kinase activity of IPMK is a well-characterized node connecting inositol polyphosphate to lipid phosphoinositide signaling, and is critical for AKT activation^3^, Wnt signaling^4^, PLC-mediated IP_3_/Calcium signaling^1,5–7^, nuclear receptor transcriptional activation^8^, mRNA export^9^, and many other roles^2,10^. However, IPMK also has enzyme-independent roles, which utilize protein-protein interactions to regulate several crucial signaling proteins. IPMK acts in an enzyme-independent manner interacting with the p53 tumor suppressor, acting as a transcriptional co-activator for p53^11,12^. IPMK participates in mTOR signaling by serving as a classic adaptor protein, connecting mTOR and Raptor in mammalian cells^13^. IPMK has been shown to mediate LKB1 regulation of AMPK by directly interacting with AMPK, which prevents AMPK phosphorylation and activation in response to several metabolic signals, including glucose^14,15^ and metformin^16^. Further, IPMK binds and stabilizes the TRAF6 ubiquitin ligase signaling scaffold in macrophages, preventing ubiquitination of the TRAF6 protein to maintain signaling^17^. Thus, these important studies have established several roles IPMK plays via kinase-dependent and–independent mechanisms.

IPMK non-catalytic functions are mediated in part by regions of the IPMK protein that are not resolved in all crystal structures of IPMK orthologues and homologues solved to date^18–23^. Although it is well established that disordered protein domains generally participate in protein-protein interactions^24–27^, what role the IPMK disordered regions might play in modulating IPMK kinase activity has not been thoroughly explored^28^.

Here, a highly engineered human IPMK lacking its two disordered domains was designed, and designated as ΔIPMK. To determine if the absence of these disordered domains alters IPMK enzyme activity on the well-known substrate PIP_2_, detailed kinetics analyses were performed and compared to wildtype activity levels. Surprisingly, ΔIPMK has a 1.8-fold higher turnover number (*k*_cat_) on PIP_2_ than wild type IPMK. In addition, a ΔIPMK construct with an additional 33 N-terminal amino acid extension to G37 (_ext_ΔIPMK) also has a 1.8-fold increase in *k*_cat_, suggesting this increase in activity is due to replacement of the native internal loop, not the unstructured N-terminus. Further, removal of the disordered domains from IPMK does not dramatically alter IPMK structure, as a 2.5 Å crystal structure of ΔIPMK maintains all the consensus features of the IPK-superfamily fold. Comparison of the ΔIPMK structure with other superfamily structures suggested a short N-terminal sequence of IPMK that is disordered in most structures could participate in ATP-nucleotide binding. This sequence was removed from ΔIPMK, and the *K*_M_ for ATP increased 4.9-fold for ΔIPMK compared to wild type IPMK. In addition, adding back a portion of the N-terminus containing the ATP clamp partially rescued the increase in *K*_M_ for ATP. Taken together, these data suggest the disordered domains of IPMK play an important role in modulating the *in vitro* kinase activity of IPMK, through multiple kinetic mechanisms.

## RESULTS

### Human IPMK lacking native internal loop is a more efficient PI3-kinase than wild type

All crystal structures of IPK-superfamily kinases have a consensus ATP-grasp fold structurally conserved between family members^18–23^. However, there are two intrinsically disordered domains in several members of the superfamily: an N-terminal disordered domain and an “internal loop” within the kinase-domain that occurs between S279 and Q373 in human IPMK (**Fig 1A**). This kinase-domain internal loop contains a nuclear localization signal and a CK2 phosphorylation site^29^. Both the N-terminal and internal loop disordered domains participate in IPMK interactions with p53^12^, TRAF6^17^, AMPK^14^, and Raptor^13^ in GST-pull down and/or functional assays, to various degrees and levels of specificity, and are usually required but not sufficient for full IPMK non-catalytic functions. Although the disordered domain’s non-catalytic functions are well established, it is unclear if they modulate IPMK kinase activity.

**Figure 1.**
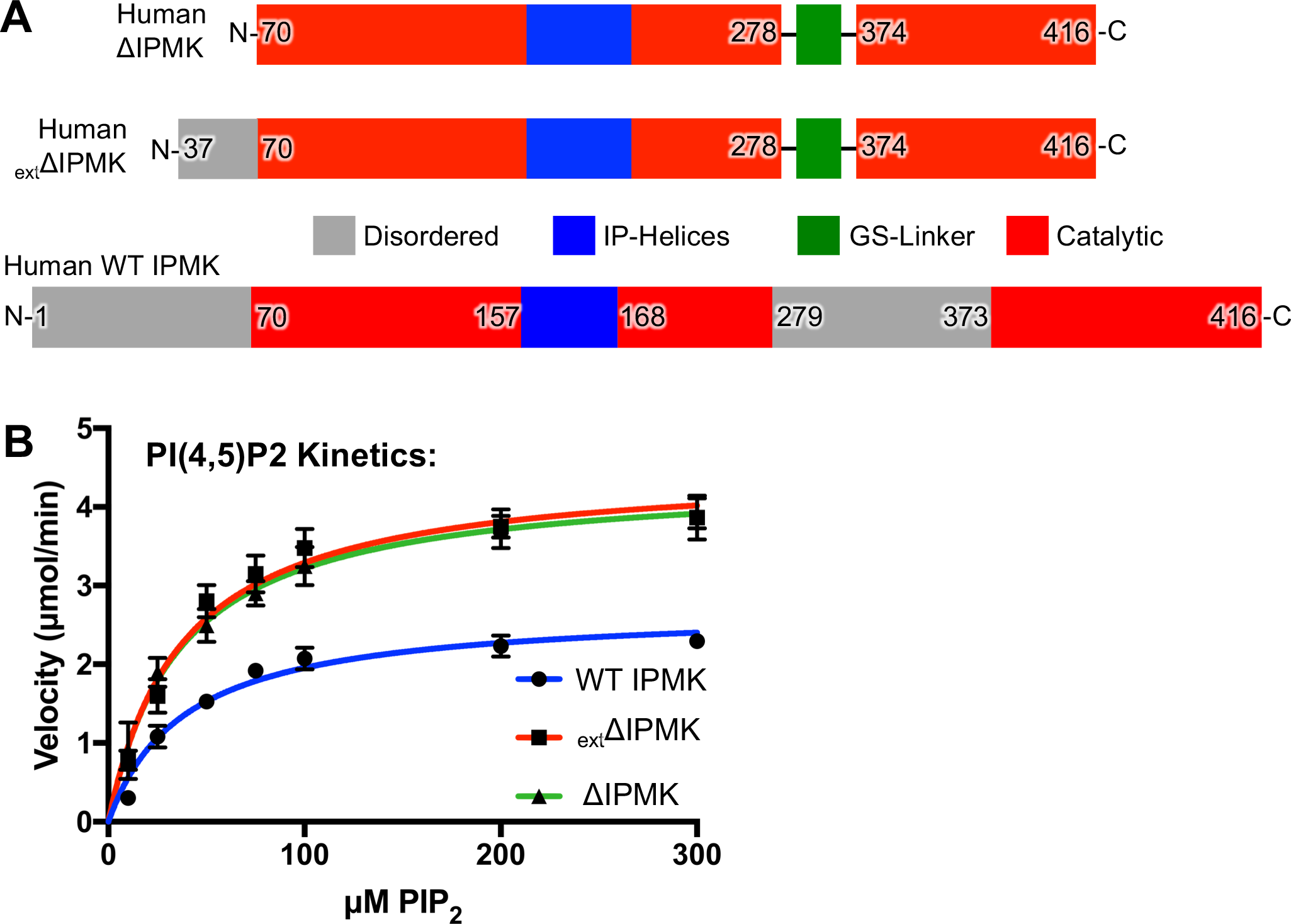
Removal of IPMK disordered domains confers a 1.8-fold increase in PIP_2_ catalytic efficiency. **A.** Primary organization comparison between human ΔIPMK (top), _ext_ΔIPMK (middle), and wild type IPMK (bottom). Gray represents disordered regions in all previous structures of IPK superfamily members, ΔIPMK does not possess residues 1-69 and 279-373 of the full-length WT human IPMK; _ext_ΔIPMK does not possess residues 1-36 and 279-373 of full length WT IPMK. Red represents catalytic regions, blue the IP-helices that bind substrate and green an artificial linker sequence added to maintain protein stability. The green artificial (Gly_4_-Ser)_2_ linker was inserted between residues 279 and 373. **B.** Best fit non-linear Michaelis-Menten curves describing WT IPMK, _ext_ΔIPMK, and ΔIPMK kinetic parameters on PIP_2_ micelles with saturating ATP, actual values provided in Table 1, 10nM of each enzyme was used in these assays. Velocities were fit using Graphpad prism.

To examine if removal of the two IPMK disordered domains alters IPMK kinetic properties on the PIP_2_ substrate, the entire disordered N-terminus of human IPMK at D70 with a 6X Histidine tag was replaced, and the kinase-domain internal loop was replaced with a small synthetic Glycine-Serine linker [S279-(GGGGS)_2_-C373] (**Fig 1A**, **S1**). This construct was designated ΔIPMK, and was easily expressed in bacteria and purified to homogeneity for crystallographic and enzyme kinetic analyses. Further, a ΔIPMK construct with an additional 33 N-terminal amino acid extension to G37 (_ext_ΔIPMK) was similarly purified and examined. It is worth noting several variations of the boundaries for this precise IPMK construct yielded significantly less amounts of soluble IPMK protein when expressed in bacteria (**Table S1**).

These kinetic studies showed although ΔIPMK has an almost identical *K*_M_ for PIP_2_ as the wild type enzyme (both about ~38 μM), the *k*_cat_ increases 1.8-fold for ΔIPMK to 7.4 s^−1^ (**Table 1**) When compared to the wild type IPMK *k*_cat_ for PIP_2_ (4.4 s^−1^), the increased *k*_cat_ drives a 1.8-fold higher catalytic efficiency (*k*_cat_/*K*_M_) of ΔIPMK over wild type IPMK (**Fig 1B**, **Table 1**). The _ext_ΔIPMK construct behaved almost identical to ΔIPMK with respect to the PIP_2_ substrate in these assays. Thus, both ΔIPMK and _ext_ΔIPMK (**Fig 1A**) appear to be more catalytically efficient than wild type IPMK at phosphorylating PIP_2_ to PIP_3_.

**Table 1:** Kinetic parameters of WT IPMK, _ext_ΔIPMK, and ΔIPMK. Graphical representations and precise boundaries of each construct are provided in Figure 1. Means ± SE using membrane capture assay, as described in Methods section.

| Kinase | $k_{\text{cat}}$ ($\text{s}^{-1}$) | $K_{\text{M,ATP}}$ ($\mu\text{M}$) | $k_{\text{cat}}/K_{\text{M,ATP}}$ ($\text{M}^{-1}\text{s}^{-1}$) | ATP $k_{\text{cat}}/K_{\text{M}}$<br>Change<br>Relative to WT | $K_{\text{M,PIP2}}$ ($\mu\text{M}$) | $k_{\text{cat}}/K_{\text{M,PIP2}}$ ($\text{M}^{-1}\text{s}^{-1}$) | PIP<sub>2</sub> $k_{\text{cat}}/K_{\text{M}}$<br>Change<br>Relative to WT |
| --- | --- | --- | --- | --- | --- | --- | --- |
| WT IPMK | $4.4 \pm 0.2$ | $61 \pm 6$ | $7.1 \times 10^4$ | 1.0 | $39 \pm 5$ | $1.2 \times 10^5$ | 1.0 |
| $\Delta_{\text{ext}}$IPMK | $7.5 \pm 0.3$ | $94 \pm 11$ | $8.0 \times 10^4$ | 1.1 | $37 \pm 5$ | $2.0 \times 10^5$ | 1.8 |
| ΔIPMK | $7.4 \pm 0.8$ | $300 \pm 64$ | $2.5 \times 10^4$ | 0.4 | $36 \pm 4$ | $2.0 \times 10^5$ | 1.8 |

### Crystallography shows ΔIPMK maintains the ATP-grasp fold

In parallel with the kinetic studies, crystals of the ΔIPMK protein were produced for X-ray structure determination of the enzyme. The growth of these crystals required the presence of the ATP-analog adenylyl-imidodiphosphate (AMP-PNP), inositol-(1,3,4,6)-phosphate (IP_4_) substrate and 1mM MnCl_2_, although these ligands were not detectable in the electron density maps. The ligands may have been lost in the 30-minute cryo-protectant soak prior to cryopreservation for X-ray diffraction analysis. Fifteen molecules of buffer components were ordered in the structure. Three of the 6 amino acids of the His-tag in ΔIPMK were weakly ordered in a position next to the β1 strand where nucleotide is predicted to be, if the nucleotide were not lost during cryopreservation. Nevertheless, we were able to solve the structure of human apo-ΔIPMK by molecular replacement using the yeast IPMK as the search model. **Table 2**contains data collection and structure refinement statistics.

**Table 2.**
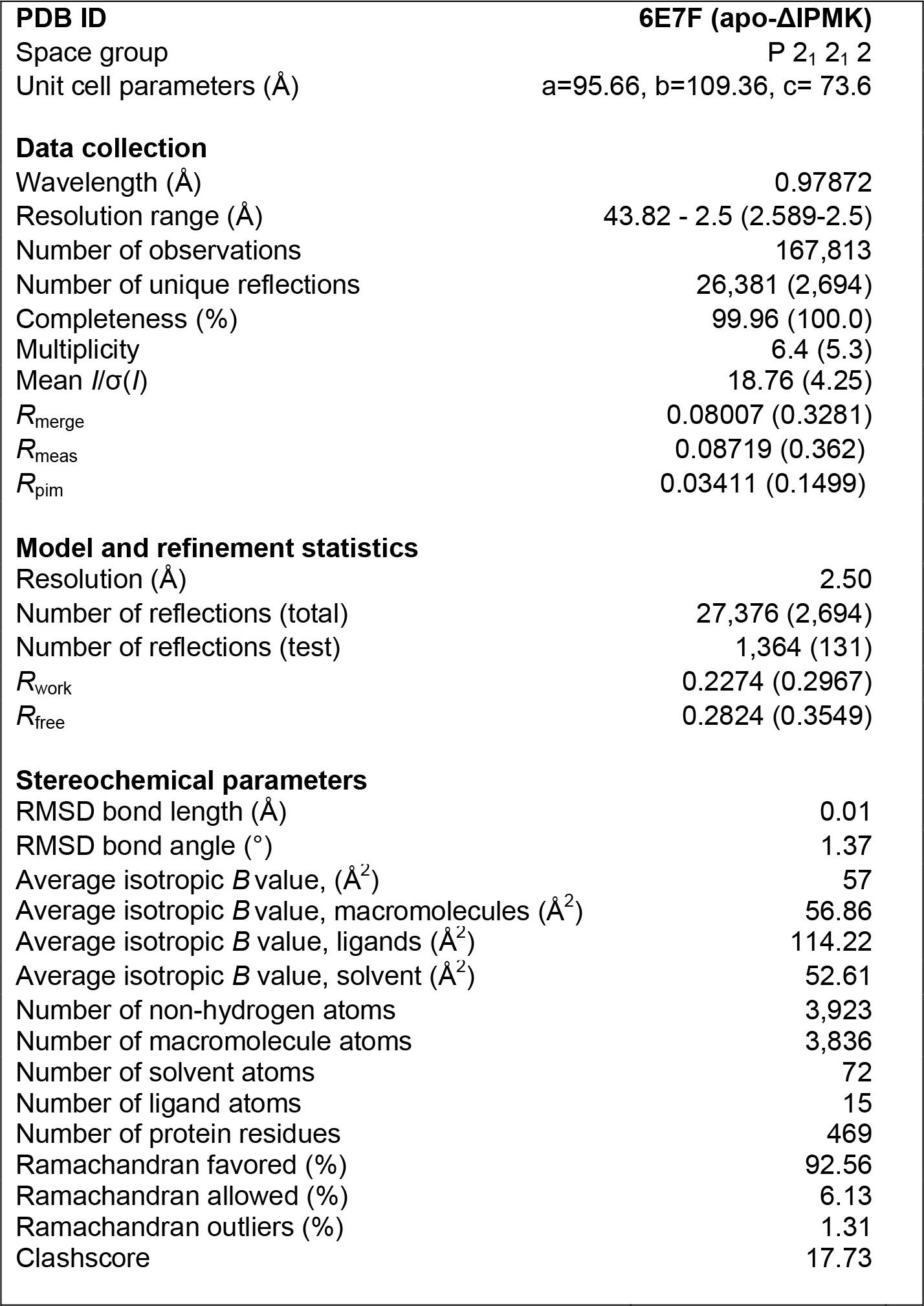
Summary of crystal parameters, data collection and refinement statistics for PDB entry 6E7F. Outermost shell shown is shown in parentheses.

|  |  |
| --- | --- |
| **PDB ID** | **6E7F (apo-ΔIPMK)** |
| Space group | P 2<sub>1</sub> 2<sub>1</sub> 2 |
| Unit cell parameters (Å) | a=95.66, b=109.36, c= 73.6 |
| **Data collection** |  |
| Wavelength (Å) | 0.97872 |
| Resolution range (Å) | 43.82 - 2.5 (2.589-2.5) |
| Number of observations | 167,813 |
| Number of unique reflections | 26,381 (2,694) |
| Completeness (%) | 99.96 (100.0) |
| Multiplicity | 6.4 (5.3) |
| Mean $I/\sigma(I)$ | 18.76 (4.25) |
| $R_{\text{merge}}$ | 0.08007 (0.3281) |
| $R_{\text{meas}}$ | 0.08719 (0.362) |
| $R_{\text{pim}}$ | 0.03411 (0.1499) |
| **Model and refinement statistics** |  |
| Resolution (Å) | 2.50 |
| Number of reflections (total) | 27,376 (2,694) |
| Number of reflections (test) | 1,364 (131) |
| $R_{\text{work}}$ | 0.2274 (0.2967) |
| $R_{\text{free}}$ | 0.2824 (0.3549) |
| **Stereochemical parameters** |  |
| RMSD bond length (Å) | 0.01 |
| RMSD bond angle (°) | 1.37 |
| Average isotropic $B$ value, (Å<sup>2</sup>) | 57 |
| Average isotropic $B$ value, macromolecules (Å<sup>2</sup>) | 56.86 |
| Average isotropic $B$ value, ligands (Å<sup>2</sup>) | 114.22 |
| Average isotropic $B$ value, solvent (Å<sup>2</sup>) | 52.61 |
| Number of non-hydrogen atoms | 3,923 |
| Number of macromolecule atoms | 3,836 |
| Number of solvent atoms | 72 |
| Number of ligand atoms | 15 |
| Number of protein residues | 469 |
| Ramachandran favored (%) | 92.56 |
| Ramachandran allowed (%) | 6.13 |
| Ramachandran outliers (%) | 1.31 |
| Clashscore | 17.73 |

Analysis of the fold of ΔIPMK revealed the expected ATP-grasp domain structure (**Fig 2A**). As with all other members of this superfamily, ΔIPMK has an N-terminal domain consisting of β-sheets 1-3 and α-helix 1, the IP-binding loop consisting of α-helix 2, and a C-terminal domain consisting of the remaining β-sheets 4-7 and α-helices 1, 3, 4, 5 and 6 (**Fig 2B**). Two ΔIPMK monomers are in each asymmetric unit of the P2_1_2_1_2 space group crystals, with a crystallographic dimerization interface mediated mainly through contacts in the IP-binding helix (**Fig 2C**). The residues participating in this interface are I92, V96, P100 and L101 (**Fig 2D**). We have not observed ΔIPMK to exist as a dimer in size exclusion chromatography, suggesting the dimer in our asymmetric unit is crystallographic. The coordinates for this structure have been deposited within the Protein Data Bank (PDB ID: 6E7F).

**Figure 2.**
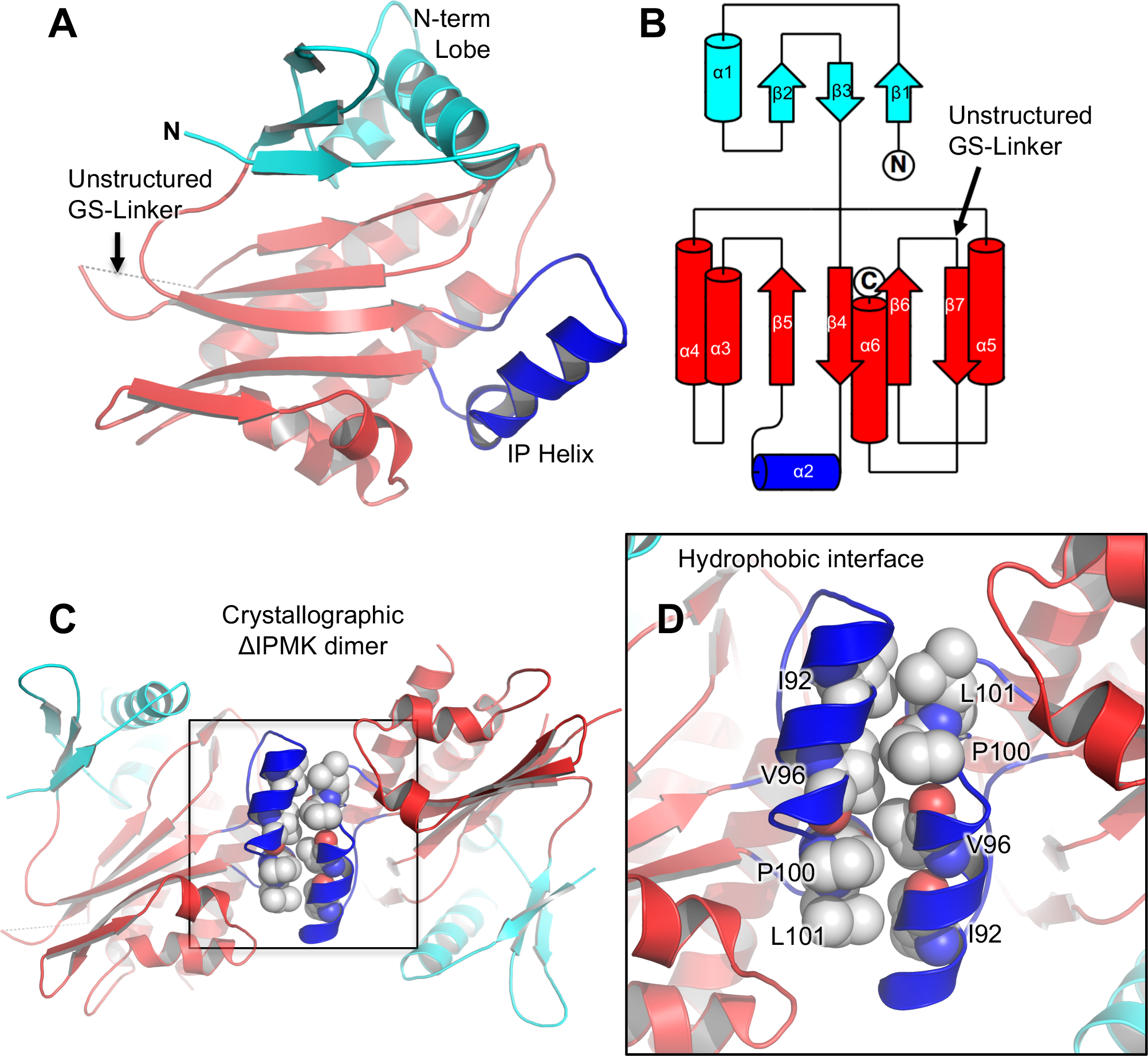
Crystal structure of the catalytic core of Human ΔIPMK. **A.** Ribbon diagrams of the overall structure of the catalytic core of human ΔIPMK and the unstructured kinase-domain internal loop shown in red, N-terminal kinase lobe shown in cyan, inositol phosphate-binding helix shown in blue. **B.** Topology map of IPMK, color scheme identical as in A., generated using TopDraw. **C.** Overall structure of the crystallographic IPMK dimer in the asymmetric unit, with the dimerization interface boxed, note that ΔIPMK dimerization is not detectable by size exclusion chromatography. **D.** Magnification of panel C, showing hydrophobic residues that mediate the crystallographic dimer interface along the IP helix, hydrophobic residues indicated.

### ΔIPMK core-kinase structure is similar to other IPK-superfamily members

Comparing ΔIPMK to other IPK-superfamily members, ΔIPMK shares an expected high degree of structural homology to other superfamily members. The structure reported here is in close agreement with the other recently published crystal structures of human IPMK, with a root mean square deviation (RMSD) of 0.48 Å^18^. Superposition over the Cα backbone of ΔIPMK with the human *H. Sapiens* IP3K gave an RMSD of 1.041 Å (**Fig 3A**). As expected, human IP3K diverges most significantly with ΔIPMK in the IP-binding loop, where IP3K has a much larger sequence that contains 2 alpha helices that are not seen in IPMK (**Fig 3A**, boxed helix). Superpositions of the human ΔIPMK with the parasitic amoeba *E. histolytica* IP6K produced an RMSD of 1.145 Å (**Fig 3B**), with plant *A. thaliana* IPMK produced an RMSD of 1.028 Å (**Fig 3C**) and with the yeast *S. cerevisiae* IPMK structure yields an RMSD of 1.10Å (**Fig 3D**). A more thorough comparison of the differences between these structures has been extensively addressed elsewhere^18^, however we note the larger IP-binding loops of IP3Ks have been proposed to prevent IP3K activity on membrane phosphoinositides^23^, and our ΔIPMK structure further supports that hypothesis.

**Figure 3.**
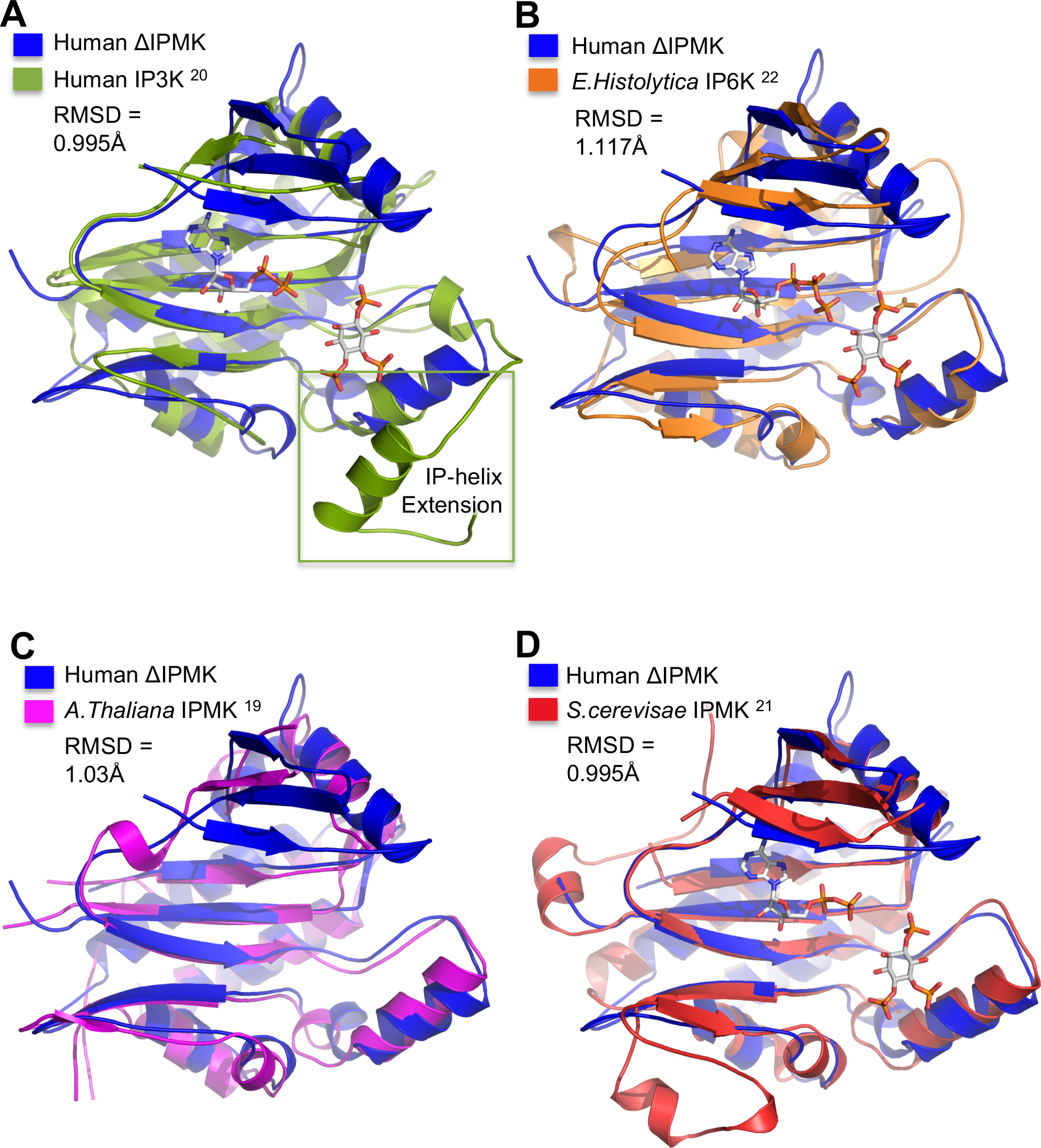
The overall fold of ΔIPMK is conserved within the Inositol Phosphate Kinase Superfamily. **A.** Superposition of ΔIPMK (blue) and human IP3K (dark green) bound to AMPPNP, Mn^2+^, and Ins(1,4,5)P_3_ (PDB: 1W2C) with a RMSD of 0.995 Å, determined using PyMOL. Human IP3K has a significantly larger inositol phosphate-binding region compared to ΔIPMK (green box), which has been ascribed to preventing IP3K from phosphorylating PIP_2_ in membranes. **B.** Superposition of ΔIPMK (blue) and *E. histolytica* IP6K (orange) bound to ATP and Ins(1,4,5)P_3_ (PDB: 4O4D) with an RMSD of 1.170Å. **C.** Superposition of ΔIPMK (blue) and apo-*A. thaliana* IPMK (magenta) (PDB: 4FRF) with an RMSD of 1.045Å. **D.** Superposition of ΔIPMK (blue) and *S. cerevisiae* IPMK (red) (PDB: 2IF8) with an RMSD of 0.995 Å. All structures generated using PyMOL and any internal loop regions disordered theses structures are represented by dashed lines connecting to the next ordered amino acid.

### ΔIPMK has higher K_M_ for ATP

In comparing our structure to other IPK-superfamily members, we noted two IPK-superfamily members contain extended N-terminal regions that are ordered in two of the crystal structures, and covers part of the nucleotide-binding pocket, as highlighted in the human IP3K structure (**Fig 4A,B**). This region is disordered in all other IPK-superfamily members except in two recently reported structures of human IPMK (PDB:5W2H and 5W2I), where a homologous region of the N-terminus, residues I65-P69 in human IPMK, is also similarly ordered over the nucleotide-binding pocket (**Fig 4C,D**) ^18^. We found the N-terminal I65 residue is in reasonable Van der Waals contact distance to the 4’ and 5’ carbons of the ribose unit of ADP (4.2Å – 3.8Å) (**Fig 4C**), but the equivalent sequence in the ΔIPMK construct was removed and replaced with an N-terminal histidine tag (**Fig 4D** and **S1**), further suggesting an ATP-stabilizing effect of the I65-P69 sequence. This sequence is clearly not required for catalysis on the PIP_2_ substrate, as the removed I65-P69 sequence in ΔIPMK results in a kinase with increased catalytic efficiency compared to the wild type enzyme (**Fig 1B, Table 1**). Nevertheless, in our electron density maps we observed weak electron density occupying a similar space as the I65-P69 sequence (**Fig 4D**). This density could only be attributed to 3 amino acids of the 6-histidine tag (**Fig 4D**). The weak density occupies a comparable space to the ordered I65-P69 sequences in the two other structures mentioned above (**Fig 4A-C**). Thus, we hypothesized the native N-terminal I65-P69 sequence might function to stabilize ATP binding in human IPMK. ΔIPMK would be predicted to have an increased *K*_M_ for ATP compared to wild type, since the I65-P69 sequence was removed from ΔIPMK. Indeed, ΔIPMK has a *K*_M_ for ATP of 300 μM (**Fig 4E**), while the ATP *K*_M_ for wild type IPMK is 61 μM (**Table 1**), demonstrating ΔIPMK has a 4.9-fold increased *K*_M_ for ATP than wild type, despite ΔIPMK having an identical *K*_M_ and 1.8-fold increased *k*_cat_ for the PIP_2_ kinase substrate (**Table 1**). When the I65-P69 sequence was added back to ΔIPMK using the _ext_ΔIPMK construct (**Fig 1A**) the *K*_M_ for ATP was rescued to 94 μM, only a 1.5-fold increase from the wild type IPMK (**Table 1**).

**Figure 4.**
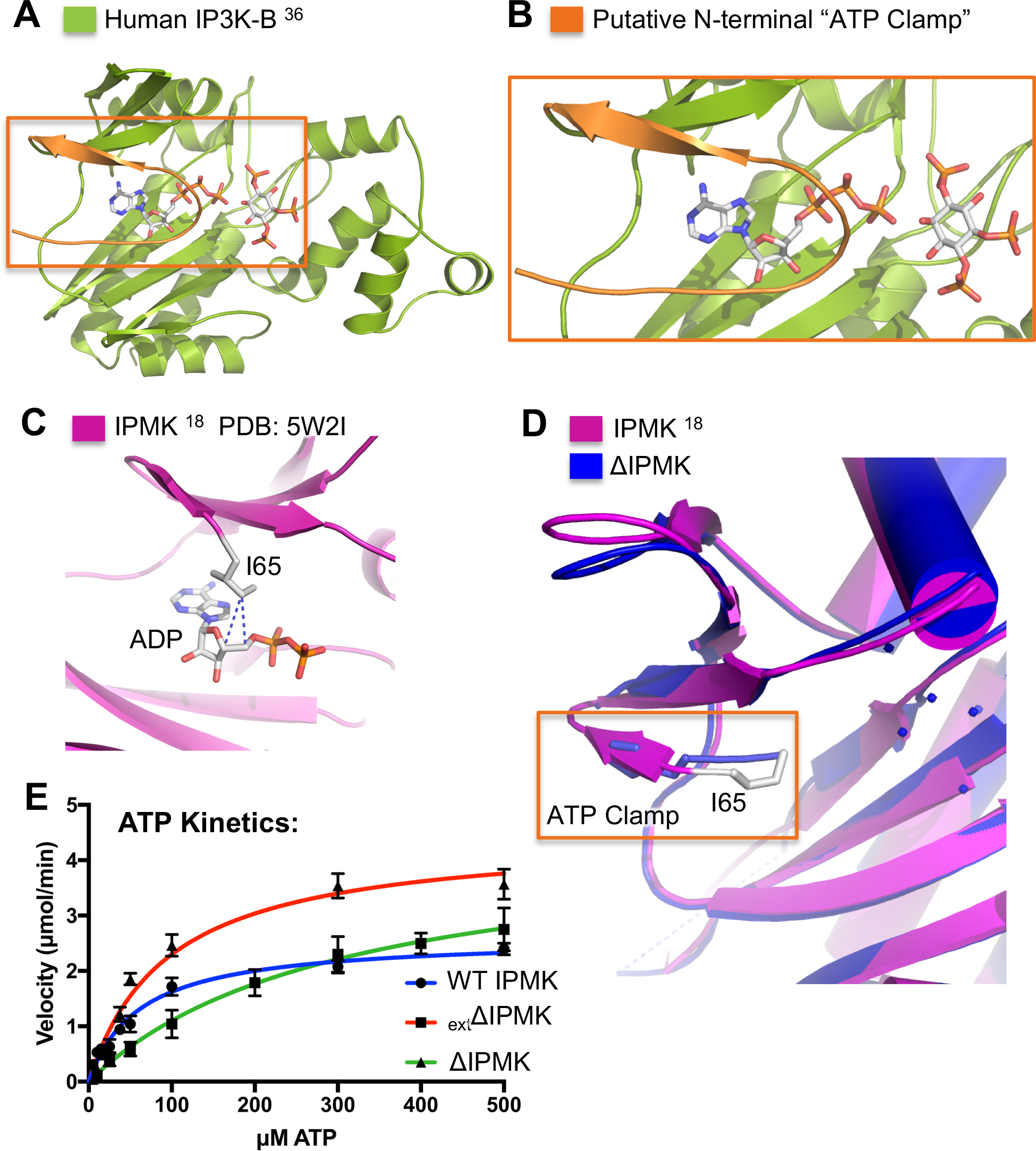
Deletion of IPMK disordered domains perturbs *K*_M_ for ATP. **A.** Ribbon diagram of IP3K-B bound to ATP with Ins(1,4,5)P_3_ modeled into the inositol binding site (PBD: 2AQX)^36^. **B.** Enlarged view of the IP3K-B ATP binding site boxed in A. shows the ordered N-terminus potentially acting as an “ATP clamp” to stabilize ATP binding. **C.** ATP binding site from the crystal structure of human IPMK bound to ADP^18^, highlighting Van der Waals interaction distances of I65 sidechain (grey sticks) to 5’ (4.2Å) and 4’ (3.8Å) carbons of the ribose moiety in ADP (sticks). **D.** N-terminus of ΔIPMK structure (blue) superposed on IPMK structure (magenta). Orange box shows the ordered ΔIPMK His-tag residues (depicted as blue sticks) in a similar position as the I65 ATP-clamp highlighted in C. **E.** Enzyme kinetic data fit to nonlinear Michaelis-Menten curves describing WT IPMK, _ext_ΔIPMK, and ΔIPMK kinetic parameters on ATP with saturating PIP_2_. Velocities were fit using Graphpad prism.

Consistent with these results, a previous study deleted the kinase-domain internal loop alone from human IPMK (residues 266-371) but left all N-terminal IPMK sequences native, and showed the *K*_M_ for ATP only increased about 2.3-fold for this mutant ^28^, suggesting removal of the kinase-domain internal loop region alone can also change the *K*_M_ for ATP. Because the change in *K*_M_ for ATP in ΔIPMK (both disordered domains removed) is 4.9-fold higher versus only a 1.5-fold change when only the kinase-domain internal loop region is removed ^28^, the data suggest changes to the N-terminal disordered domain immediately proceeding the kinase domain more dramatically determine ATP binding affinity, but also that there exists a synergy between both disordered domains which can alter ATP binding to IPMK. Taken together, these crystallographic observations and enzyme kinetic data suggest the native N-terminal sequences of human IPMK can form a previously uncharacterized and structurally conserved ATP clamp, stabilizing ATP binding to IPMK.

## DISCUSSION

Here, we report removal of the disordered domains of human IPMK (ΔIPMK) results in an enzyme that is more catalytically efficient than wild type IPMK, driven by a 1.8-fold increase in *k*_cat_. Since the *K*_M_ for PIP_2_ is not significantly different between ΔIPMK, _ext_ΔIPMK, or wild type IPMK (**Table 1**), IPMK disordered domains are not likely involved with PIP_2_ substrate binding. In support of this claim, as 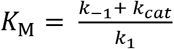, *K*_M,WT_ = *K*_M,extΔIPMK_ = *K*_M,Δ_ for PIP_2_, and *k*_cat,WT IPMK_ < *1k*_cat,extΔIPMK_ = *k*_cat,Δ_ for PIP_2_, *k*_cat_ does not substantially influence the IPMK *K*_M_ for PIP_2_. Therefore, IPMK *K*_M,PIP2_ ≈ *K*_D,PIP2_, which suggests removal of the native IPMK internal loop and replacement with a glycine-serine linker does not alter IPMK’s binding affinity for PIP_2_. If the IPMK disordered domains did affect PIP_2_ binding, the PIP_2_ *K*_M_ would be expected to change for both _ext_ΔIPMK and ΔIPMK compared to wild type IPMK. In agreement with our data, a previous study suggested completely removing the human IPMK internal loop domain (residues 266-371) does not alter substrate binding, since removal of the domain alone only nominally increased IPMK enzyme substrate *K*_M_ for the Inositol (1,4,5) trisphosphate (IP_3_) substrate (0.35 μM for WT *vs.*0.4 μM for mutant)^28^. Thus, the data clearly suggest neither of the two disordered domains in human IPMK affect substrate binding to the IP-helix.

As an alternative hypothesis, IPMK disordered domains could be predicted to alter ATP binding, even when excess ATP is present as is the case for PIP_2_ kinetics experiments. However, the *K*_M_ for ATP is actually increased 1.5-fold for _ext_ΔIPMK and 4.9-fold for ΔIPMK, despite the enzyme becoming faster. Thus, the data suggest the disordered domains actually increase IPMK binding affinity for ATP because the *K*_M_ for ATP decreases with increasing proportions of the disordered domains present (i.e. *K*_M,*WT*_ < *K*_M,*extΔ*_ ≪ *K*_M, *Δ*_).

The structure of ΔIPMK allowed us to highlight regions of IPMK previously established to be required in GST pull-down assays for full co-precipitation with p53 (**Fig 5A,B**). The p53-IPMK interaction was shown to modulate p53 activity, and p53 requires the IPMK IP-binding loop and ATP cleft to fully interact with IPMK. This model predicts interaction of p53 with IPMK would down-regulate IPMK kinase activity, consistent with p53 serving an anti-proliferative role during stress responses. Similar analyses using AMPK (**Fig S2A**) and TRAF6 (**Fig S2B**) were also consistent with protein-protein interactions inhibiting IPMK kinase activity, although this remains to be determined.

**Figure 5.**
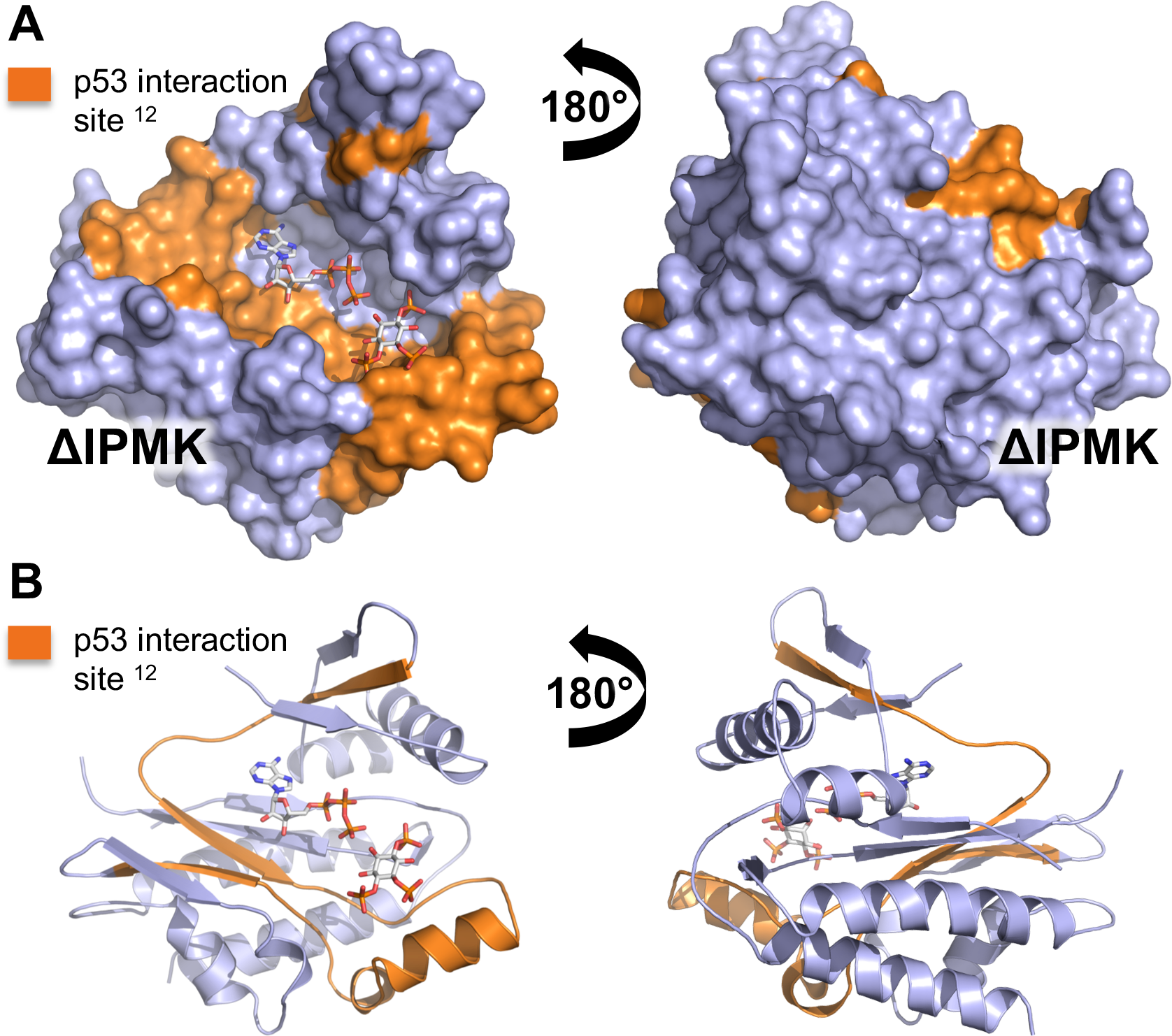
p53 interaction site superimposes onto ΔIPMK ATP- and substrate-binding sites. **A.** Surface and **B.** cartoon ribbon representations of ΔIPMK, with ADP and PIP_2_ modeled into the structure. The IPMK interaction site with p53 has been mapped to IPMK exon 4^12^, the residues of exon 4 are highlighted in orange, with the remainder of the IPMK kinase domain depicted in light blue. Both ADP and an inositol phosphate kinase substrate were modeled into the structure for reference, the position of the p53 interaction site predicts p53 could function to sterically interfere with IPMK kinase activity.

Taken together, the crystallographic and kinetic data are consistent with a model of IPMK where the disordered domains allosterically modulate kinase activity, through a mechanism that does not alter kinase substrate binding affinity (i.e. PIP_2_), increases ATP nucleotide binding affinity (i.e. decreases *K*_M_), and decreases the turnover number (i.e. *k*_cat_) (**Fig 6A,B**). This could represent biologically relevant regulation, given intrinsically disordered domains are well known to serve as protein-protein interaction scaffolding domains. Proteins that interact with IPMK disordered loops could compete for disordered domain binding, freeing IPMK from allosteric inhibition (**Fig 6C**). However, it is not clear from a structural perspective how any allosteric mechanism could work. Both disordered domains are present in the yeast IPMK ^21^ crystal structure (although no electron density is observed in these regions), and yet the fold of the yeast IPMK with disordered domains is very similar to human IPMK without the disordered domains. It remains possible that small perturbations in side chain interactions between the IPMK core kinase domain and disordered domains could regulate catalytic rate. Thus, solution-based biophysical dynamics data may be required to resolve this question, such as heteronuclear spin quantum coupling NMR studies to determine sites of contact and electron paramagnetic resonance (EPR) based double electron-electron resonance (DEER) spectroscopy measurements to determine long-range distances between residues.

**Figure 6:**
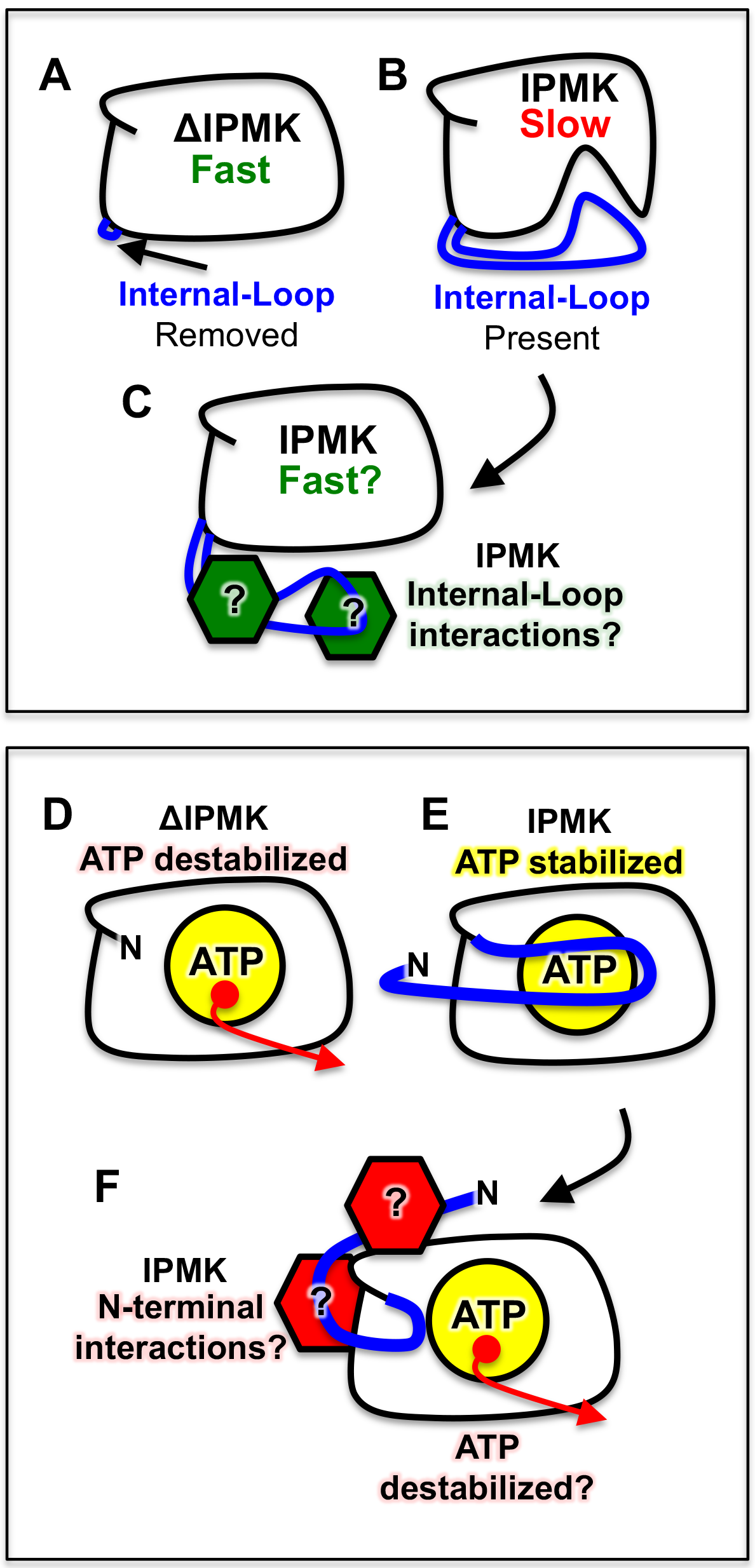
Model of intrinsic regulation of IPMK catalysis and ATP-binding. **A.** Removal of disordered domains (blue) from ΔIPMK results in a catalytically faster enzyme. **B**. When present, the disordered domains (blue) may allosterically modulate the core kinase domain of IPMK that results in a slower enzyme. **C.** Proteins (green) could interact with the internal loop region of IPMK, removing the inhibition of the disordered domains, resulting in activation of IPMK catalysis. **D.** Removal of disordered domains from ΔIPMK results in less stable ATP binding. **E.** When present, the disordered domains stabilize ATP binding through the N-terminal regions that includes the I65-P69 ATP-clamp sequence (blue). **F.** Proteins that interact with the disordered N-terminal region of IPMK (red) could regulate the structural positioning of the ATP clamp sequence, hence regulating IPMK ATP-binding in certain cellular compartments where ATP concentrations are low.

A recently published structure of apo-human IPMK had a similar but non-identical construct design as reported here^18^, and comparison of our 2.5 Å apo-ΔIPMK structure to the 1.8 Å apo-*Hs*IPMK structure (PDB: 5W2G) reveals almost identical results, with an RMSD of 0.48 Å over the 161 identical residues of the IPMK Cα backbone, despite different space groups and different numbers of molecules in the crystallographic asymmetric units. Further, the crystallization conditions reported here required acidic mother liquor (sodium citrate pH 3.5), while the conditions reported for *H.s.*IPMK are far milder (MES-imidazole pH 6.0 or HEPES pH 7.5). Despite these differences, the final structures derived from these crystals are very similar. Thus, the independently solved structures make a strong case the native structure of the human IPMK core kinase domain is accurately represented with these models, at least as examined by X-ray crystallography.

In two of the structures of human IPMK (PDB: 5W2H, 5W2I), a short 5-amino acid N-terminal sequence (I65-P69) was ordered directly above the ATP binding site^18^. These residues, particularly I65, form a “roof” atop the ATP binding pocket, with the well-resolved I65 side chain in reasonable Van der Waals distance (4.2Å – 3.8Å) from the nucleotide ribose sugar C4’ and C5’ positions (**Fig 4C**) ^18^. Similar sequences over the ATP-binding sites are ordered in the crystal structure of the IPK-superfamily members human IP3K^20,23^ and the amoeba IP6K^22^, but are disordered in all other structures. These crystallographic observations suggested residues I65-P69 of human IPMK might form an “ATP-clamp” helping to hold ATP in the nucleotide-binding pocket (**Fig 6E**), and that removal of the clamp would increase the *K*_M_ for ATP (**Fig 6D**). The I65-P69 ATP-clamp sequence was deleted from the ΔIPMK construct reported here, and accordingly observed a 4.9-fold increase in the *K*_M_ for ATP in ΔIPMK, but only a 1.5 fold increase in ATP *K*_M_ was observed when the ATP-clamp sequence was restored using the _ext_ΔIPMK construct.

It is unknown if the ATP clamp sequence can alter IPMK activity in living cells. Since intracellular ATP levels normally range between 2-10 mM^30–32^, one might expect shifting the *K*_M_ for ATP from 61 μM to 300 μM to have little if any regulatory capacity on IPMK kinase activity. However, ATP concentrations in certain subcellular compartments can exist below the level of detection for ATP biosensors with dissociation constants for ATP below 10 μM^30,33^, and ATP levels can change in subcellular compartments as measured with biosensors by over 1,000-fold^31,32,34^. IPMK shuttles between the nucleus and the cytoplasm^29^, but often localizes to the nucleus with a speckled, uneven distribution^3^. The I65-P69 sequence is not ordered in most IPK-superfamily structures, suggesting the I65-P69 sequence is dynamic. It is tempting to speculate IPMK ATP binding could be regulated by structural perturbations induced in the I65-P69 ATP clamp caused by protein-protein interactions with the N-terminal intrinsically disordered domain of IPMK (**Fig 6F**). In this model, the positioning of the I65-P69 ATP-clamp over the nucleotide could be modulated by proteins interacting with the disordered N-terminus of IPMK (**Fig 6F**). This model predicts proteins discovered to interact with the N-terminus of human IPMK might function to discourage ATP binding to IPMK, perhaps regulating IPMK kinase activity in certain sub-cellular compartments when ATP levels drop. This type of regulation is further supported by GST-pull down interaction data showing proteins require the substrate-binding helix and the ATP binding cleft to fully interact with IPMK (**Fig 5**). Another intriguing possibility is the recent elucidation that the nucleotide GTP can be used by a certain phosphoinositide kinase previously thought to only use ATP for the phosphoryl-transfer reaction^35^. Should the ATP-clamp help select for GTP or ATP, any regulation of the clamp might have the possibility to act as a “nucleotide switch”, converting IPMK from an ATP-dependent kinase to a GTP-dependent kinase. It is currently unknown if IPMK can use GTP in the phosphoryl-transfer reaction.

From a structural perspective, it is important to note all the published crystal structures of IPK-superfamily members appear to be extremely similar with or without small molecule ligands bound to the kinase, e.g. superposition of the recently published IPMK (PDB:5W2I) bound to 3 ligands (ATP-analog, 2 magnesium metal ions and diC_4_-PIP_2_) and the apo-ΔIPMK reported here has an RMSD over the Cα backbone of 0.414Å. Similarly, superposition of ΔIPMK vs. ADP/Mn^2+^-bound *S. cerevisiae* IPMK^21^ has an RMSD of 0.995Å over the Cα backbone. These data suggest the IPK-superfamily of kinases do not undergo large conformational changes upon small molecule ligand binding. However, it remains possible these ligands induce conformational dynamics less well suited to be detected by X-ray crystallography and which could become apparent if solution biophysical approaches were applied.

The crystallographic and kinetic studies presented here show the disordered domains of human IPMK can inhibit IPMK *in vitro* kinase activity on PIP_2_ not by altering PIP_2_ binding, but by somehow decreasing the turnover number. We also show that removal of the IPMK disordered domains discourages ATP binding to IPMK *in vitro*. The data are consistent with a new model where the disordered domains of IPMK modulate its kinase activity, through multiple, distinct kinetic mechanisms. Crystallographic studies confirmed removing the disordered domains does not alter the overall fold of the IPMK kinase-domain, however the structural basis of how IPMK disordered domains alter the kinase activity remains to be determined.

## METHODS

### Protein Expression and Purification

The human ΔIPMK expression vector was constructed by VectorBuilder (Santa Clara, CA). Briefly, human IPMK (NCBI nucleotide accession NM_152230.4) 70-279 and 373-416 interrupted with a (Gly_4_-Ser)_2_ linker was inserted into an IPTG inducible N-terminal His-TEV expression vector. This expression vector was transformed into chemically competent BL21 dE3 *E. coli* cells. An overnight Terrific Broth (Sigma) culture under ampicillin selection of transformed *E. coli* was inoculated into 4L of Terrific broth, grown at 37°C to OD_600_=0.8, and induced with 1mM IPTG at 15°C overnight. Cells were resuspended in Buffer A (20 mM Tris pH 8.0, 300 mM NaCl, 20mM imidazole, 5% glycerol, 5 mM βME) with protease inhibitor tablets (Roche) and frozen at −80° C until ready for use. Cells were lysed via sonication at 4°C for 10 minutes and centrifuged at 16,000×g for 20 minutes at 4°C. Supernatant was incubated with TALON^®^ metal affinity resin (Clontech) for 1 hour, washed with 20 column volumes of Buffer A, and eluted with Buffer A + 200mM imidazole. Eluted protein was loaded directly onto a SEPAX SRT-10 SEC-300 sizing column and desalted into Buffer B (20 mM Tris pH 8.0, 5% glycerol, and 5mM βME). Protein fractions were then loaded onto a MonoQ 5/50 GL anion exchange column (GE Life Sciences) using an AKTA PURE (GE Life Sciences) and eluted using a gradient of Buffer B + 10mM NaCl. ΔIPMK peaks were concentrated to 10mg/mL and used for both enzyme assays and crystallization experiments.

### IPMK Kinase Assays

IPMK activity on PI(4,5)P_2_ (PIP_2_) kinase assays were performed using radiolabeled nitrocellulose capture assays as previously described^8^. In all kinetics experiments, lyophilized dipalmitoyl PIP_2_ under a vacuum was resuspended in water and sonicated with dioleyl-phosphatidylserine (DOPS) to form micelles. The final concentration of all enzymes were 10 nM, added to reactions with *in vitro* kinase buffer (50 mM HEPES pH 7.5, 5 mM MgCl_2_). Cold ATP was used at 10 mM final concentration to saturate the enzyme with respect to ATP when measuring PIP_2_ kinetics, and 300 μM PIP_2_ was used to saturate IPMK with respect to PIP_2_ when determining ATP kinetics. Cold ATP was spiked with purified 0.25 μCi/μL [^32^P]γ-ATP (Perkin Elmer) in reactions. All reactions were initiated by addition of ATP, incubated at 37°C for one minute, and quenched by dotting onto nitrocellulose paper. One minute was determined empirically to maximize signal/noise while remaining within the steady state of the reaction. Radioactivity incorporation was detected using a phosphorimaging screen, scanned on a Typhoon phosphoimager and quantified using ImageJ. Data were processed using a standard Michaelis-Menten curve fit using GraphPad Prism.

### Crystallization

ΔIPMK crystallization conditions were screened with the Joint Center for Structural Genomics (JCSG) Core Suite using the TTP Labtech Mosquito^®^ with 100nL drops via sitting drop vapor diffusion at a final protein concentration of 3.33 mg/mL. Crystals formed after 3 days in 0.1M sodium citrate pH 3.5, 0.8M (NH_4_)_2_SO_4_ in the presence of 1 mM AMPPNP, 1 mM MnCl_2_, and 0.73 mM Ins(1,3,6)P_4_. Crystals were cryo-protected in crystallization buffer + 25% ethylene glycol for 30 minutes, and flash frozen in liquid nitrogen for X-ray diffraction data collection.

### Crystal Data Collection, Structure Determination, and Refinement

Diffraction data were collected using APS beamline 21-ID-F and processed with HKL2000. Initial phases were determined using the BALBES automated molecular replacement pipeline, which empirically determined the *S. cerevisiae* IPMK bound to ADP (PDB: 2IF8) as the highest confidence search model. The initial structure was rebuilt using COOT and refined with phenix.refine in the Phenix software suite. Structural model figures were generated with Pymol and UCSF Chimera. Of note, 6 Ramachandran outliers were observed. Two of these 6 were part of the artificial and very poorly ordered His-tag (Chain A: H-13, and Chain B: H-13). One of the remaining 4 outliers is a native IPMK amino acid (Chain B: S261), however S261 is the second to the last assignable residue preceding the highly disordered and unassignable Ser/Glycine linker region, itself in a region of low density. Inclusion of the artificial Ser/Gly linker in the construct was required for crystallization, as also noted by others^18^. The remaining 3 Ramachandran outliers are all native IPMK amino acids (Chain A: S188; Chain B: K136, C220, S261), are also most likely due to poor electron density in these three areas of the model. None of the outlier amino acids are in an obvious crystallographic interface between asymmetric units, or in any interface between Chains A and B. Structure factors and atomic coordinates were deposited in the Protein Data Bank (PDB) with accession code **PDB: 6E7F**.

### Data availability

The datasets generated and analysed during the current study are available in the Protein Data Bank repository as 6E7F, at https://www.rcsb.org/.

## Author Contributions

CDS performed all experiments, CDS and RDB designed and interpreted data from all the experiments, wrote the manuscript text, prepared all figures and reviewed the final manuscript text.

## Supplementary Information

**Figure S1.**
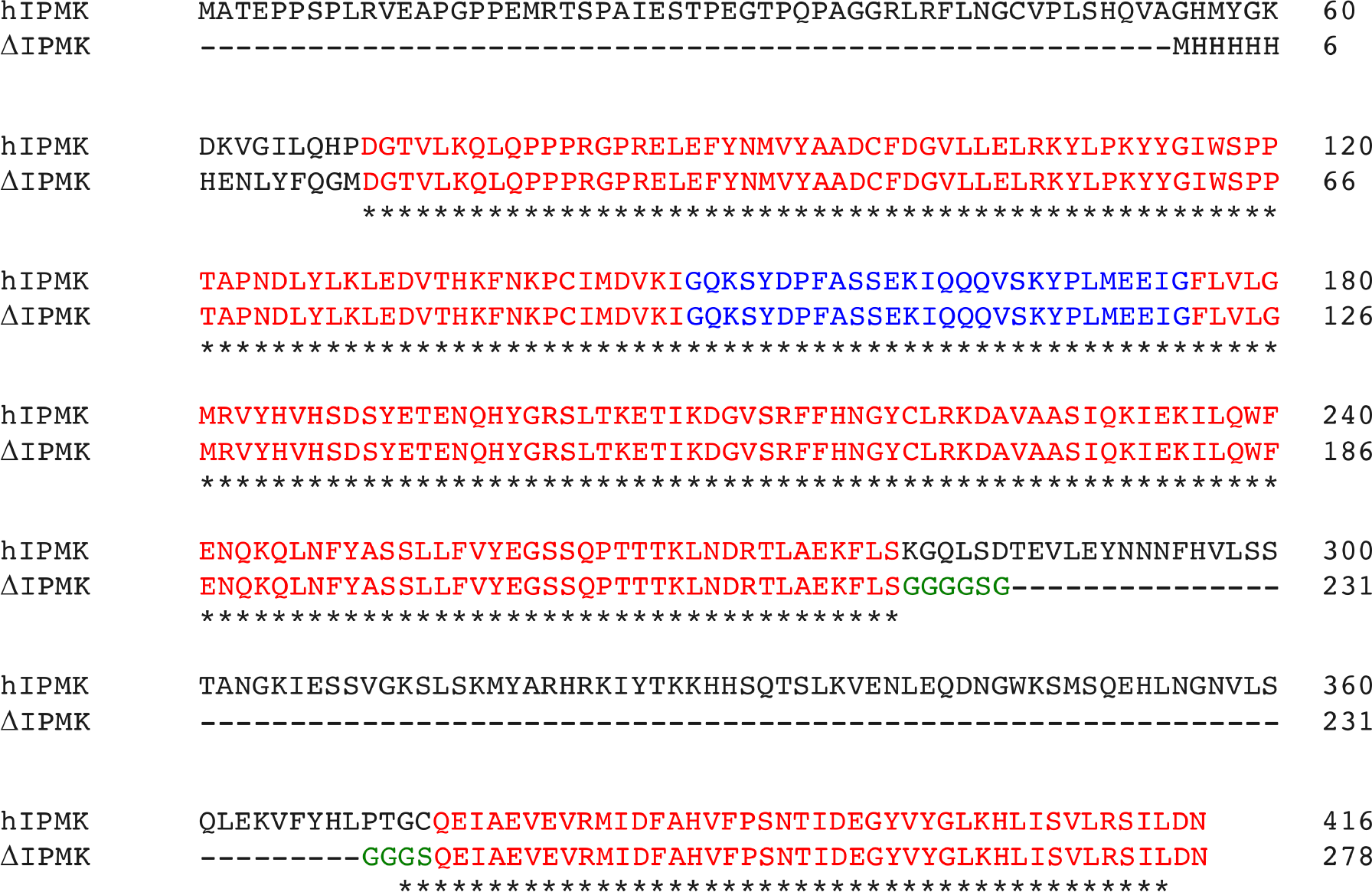
Sequence alignment of full length human IPMK and ΔIPMK. ΔIPMK does not possess residues 1-69 and 279-373 of the full-length human IPMK sequence. Red represents catalytic regions, blue the IP-helices that bind substrate and green an artificial linker sequence added to maintain protein stability. The green artificial (Gly_4_-Ser)_2_ linker was inserted between residues 279 and 373.

**Table S1.** Human IPMK construct expression in bacteria and crystallization.

| N-term Domain Boundary | Internal-Loop Boundaries | Tags | Internal-Loop Linker | Relative Expression | Crystals |
| --- | --- | --- | --- | --- | --- |
| D70 | S279-C373 | N-term 6XHis | (GGGGS)2 | +++ | Yes |
| G37 | S279-C373 | C-term 6XHis | (GGGGS)2 | + | None |
| M1 | Q262-P370 | N-term 6XHis-MBP | None | - | n.d. |
| G59 | Q262-P370 | N-term 6XHis MBP | None | - | n.d. |

**Figure S2.**
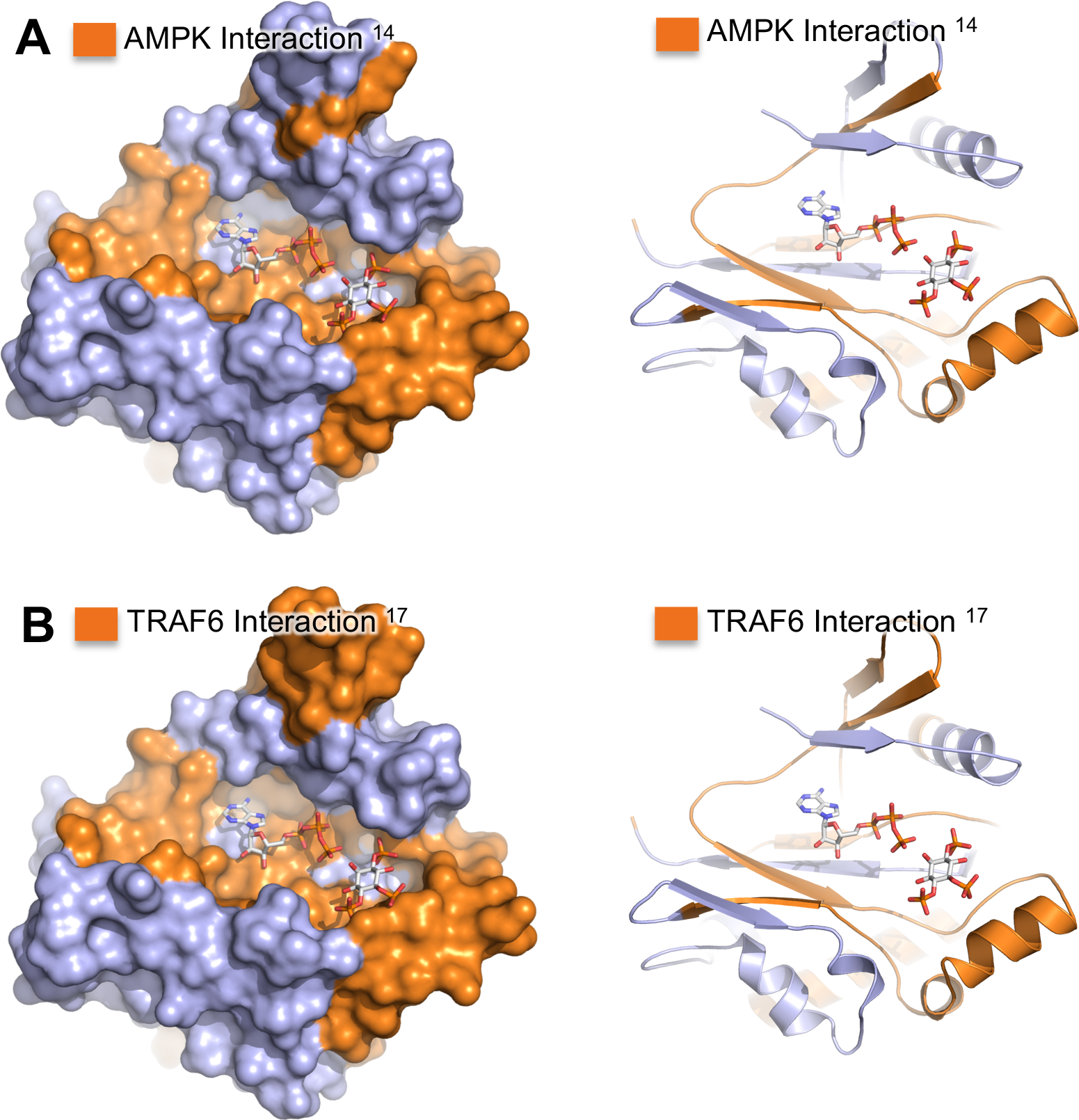
Protein-protein interaction sites superimpose onto IPMK ATP- and substrate-binding sites. Surface (left) and ribbon (right) representations of ΔIPMK, with ADP and PIP_2_ modeled into the structure. **A.** IPMK interaction site with AMPK has been mapped^14^ to IPMK exons 4 and 6, depicted as orange. With the remainder of the IPMK kinase domain is depicted in light blue. Both ADP and an inositol phosphate kinase substrate were modeled into the structure for reference. **C.** IPMK interaction site with TRAF6 has been mapped^17^ to exons 4 and 6, depicted as orange. Note that in both cases, binding of these proteins to IPMK would be predicted to sterically interfere with IPMK kinase activity.

## REFERENCES

1. Frederick, J. P. et al. An essential role for an inositol polyphosphate multikinase, Ipk2, in mouse embryogenesis and second messenger production. Proc Natl Acad Sci U S A 102, 8454–8459 (2005).

2. Kim, E., Beon, J., Lee, S., Park, J. & Kim, S. IPMK: A versatile regulator of nuclear signaling events. Adv. Biol. Regul. 61, 25–32 (2016).

3. Resnick, A. C. et al. Inositol polyphosphate multikinase is a nuclear PI3-kinase with transcriptional regulatory activity. Proc Natl Acad Sci U S A 102, 12783–12788 (2005).

4. Gao, Y. & Wang, H. Inositol pentakisphosphate mediates Wnt/beta-catenin signaling. J. Biol. Chem. 282, 26490–502 (2007).

5. York, S. J., Armbruster, B. N., Greenwell, P., Petes, T. D. & York, J. D. Inositol diphosphate signaling regulates telomere length. J. Biol. Chem. 280, 4264–9 (2005).

6. Leyman, A. et al. The absence of expression of the three isoenzymes of the inositol 1,4,5-trisphosphate 3-kinase does not prevent the formation of inositol pentakisphosphate and hexakisphosphate in mouse embryonic fibroblasts. Cell. Signal. 19, 1497–504 (2007).

7. Fujii, M. & York, J. D. A Role for Rat Inositol Polyphosphate Kinases rIPK2 and rIPK1 in Inositol Pentakisphosphate and Inositol Hexakisphosphate Production in Rat-1 Cells. J. Biol. Chem. 280, 1156–1164 (2005).

8. Blind, R. D., Suzawa, M. & Ingraham, H. A. Direct Modification and Activation of a Nuclear Receptor-PIP2 Complex by the Inositol Lipid Kinase IPMK. Sci Signal 5, ra44 (2012).

9. Wickramasinghe, V. O. et al. Human inositol polyphosphate multikinase regulates transcript-selective nuclear mRNA export to preserve genome integrity. Mol Cell 51, 737–750 (2013).

10. Malabanan, M. M. & Blind, R. D. Inositol polyphosphate multikinase (IPMK) in transcriptional regulation and nuclear inositide metabolism. Biochem. Soc. Trans. 44, (2016).

11. Xu, R. & Snyder, S. H. Gene transcription by p53 requires inositol polyphosphate multikinase as a co-activator. Cell Cycle 12, 1819–1820 (2013).

12. Xu, R. et al. Inositol polyphosphate multikinase is a coactivator of p53-mediated transcription and cell death. Sci Signal 6, ra22 (2013).

13. Kim, S.-Y. et al. Amino Acid Signaling to mTOR Mediated by Inositol Polyphosphate Multikinase. Cell Metab. 13, 215–221 (2011).

14. Bang, S. et al. AMP-activated protein kinase is physiologically regulated by inositol polyphosphate multikinase. Proc Natl Acad Sci U S A 109, 616–620 (2012).

15. Saiardi, A., Erdjument-Bromage, H., Snowman, A. M., Tempst, P. & Snyder, S. H. Synthesis of diphosphoinositol pentakisphosphate by a newly identified family of higher inositol polyphosphate kinases. Curr. Biol. 9, 1323–1326 (1999).

16. Bang, S., Chen, Y., Ahima, R. S. & Kim, S. F. Convergence of IPMK and LKB1-AMPK signaling pathways on metformin action. Mol. Endocrinol. 28, 1186–1193, 8 pp (2014).

17. Kim, E. et al. Inositol polyphosphate multikinase promotes Toll-like receptor-induced inflammation by stabilizing TRAF6. Sci. Adv. 3, e1602296 (2017).

18. Wang, H. & Shears, S. B. Structural features of human inositol phosphate multikinase rationalize its inositol phosphate kinase and phosphoinositide 3-kinase activities. J. Biol. Chem. 292, 18192–18202 (2017).

19. Endo-Streeter, S., Tsui, M. K., Odom, A. R., Block, J. & York, J. D. Structural studies and protein engineering of inositol phosphate multikinase. J Biol Chem 287, 35360–35369 (2012).

20. Miller, G. J. & Hurley, J. H. Crystal structure of the catalytic core of inositol 1,4,5- trisphosphate 3-kinase. Mol. Cell 15, 703–11 (2004).

21. Holmes, W. & Jogl, G. Crystal structure of inositol phosphate multikinase 2 and implications for substrate specificity. J Biol Chem 281, 38109–38116 (2006).

22. Wang, H., DeRose, E. F., London, R. E. & Shears, S. B. IP6K structure and the molecular determinants of catalytic specificity in an inositol phosphate kinase family. Nat. Commun. 5, 4178 (2014).

23. González, B. et al. Structure of a human inositol 1,4,5-trisphosphate 3-kinase: substrate binding reveals why it is not a phosphoinositide 3-kinase. Mol. Cell 15, 689–701 (2004).

24. Wright, P. E. & Dyson, H. J. Intrinsically disordered proteins in cellular signalling and regulation. Nat. Rev. Mol. Cell Biol. 16, 18–29 (2015).

25. Tompa, P. Intrinsically disordered proteins: a 10-year recap. Trends Biochem. Sci. 37, 509–516 (2012).

26. Tompa, P., Schad, E., Tantos, A. & Kalmar, L. Intrinsically disordered proteins: emerging interaction specialists. Curr. Opin. Struct. Biol. 35, 49–59 (2015).

27. Uversky, V. N. Dancing Protein Clouds: The Strange Biology and Chaotic Physics of Intrinsically Disordered Proteins. J. Biol. Chem. 291, 6681–6688 (2016).

28. Mayr, G. W., Windhorst, S. & Hillemeier, K. Antiproliferative plant and synthetic polyphenolics are specific inhibitors of vertebrate inositol-1,4,5-trisphosphate 3-kinases and inositol polyphosphate multikinase. J Biol Chem 280, 13229–13240 (2005).

29. Meyer, R. et al. Nucleocytoplasmic shuttling of human inositol phosphate multikinase is influenced by CK2 phosphorylation. Biol. Chem. 393, 149–60 (2012).

30. Yoshida, T., Alfaqaan, S., Sasaoka, N. & Imamura, H. in Methods in molecular biology (Clifton, N.J.) 1567, 231–243 (2017).

31. Suzuki, R., Hotta, K. & Oka, K. Spatiotemporal quantification of subcellular ATP levels in a single HeLa cell during changes in morphology. Sci. Rep. 5, 16874 (2015).

32. Ando, T. et al. Visualization and Measurement of ATP Levels in Living Cells Replicating Hepatitis C Virus Genome RNA. PLoS Pathog. 8, e1002561 (2012).

33. Vancraenenbroeck, R. & Webb, M. R. A Fluorescent, Reagentless Biosensor for ATP, Based on Malonyl-Coenzyme A Synthetase. ACS Chem. Biol. 10, 2650–2657 (2015).

34. Imamura, H. et al. Visualization of ATP levels inside single living cells with fluorescence resonance energy transfer-based genetically encoded indicators. at <http://www.pnas.org/content/pnas/106/37/15651.full.pdf>

35. Sumita, K. et al. The Lipid Kinase PI5P4Kβ Is an Intracellular GTP Sensor for Metabolism and Tumorigenesis. Mol. Cell 61, 187–198 (2016).

36. Chamberlain, P. P. et al. Structural insights into enzyme regulation for inositol 1,4,5-trisphosphate 3-kinase B. Biochemistry 44, 14486–93 (2005).

